# Phospholamban Inhibits the Cardiac Calcium Pump Through Reversing the Allosteric Enhancement of Calcium Affinity by ATP

**DOI:** 10.1101/2023.07.20.549928

**Authors:** Sean R. Cleary, Jaroslava Seflova, Ellen E. Cho, Konark Bisht, Himanshu Khandelia, L. Michel Espinoza-Fonseca, Seth L. Robia

**Author notes:** To Whom Correspondence Should be Addressed:* Seth L. Robia, Department of Cell and Molecular Physiology, Stritch School of Medicine, Loyola University Chicago, 2160 S. First Ave. Maywood, IL 60153.

## Abstract

Phospholamban (PLB) is a transmembrane micropeptide that regulates the Ca^2+^ pump SERCA in cardiac muscle, but the physical mechanism of this regulation remains poorly understood. PLB reduces the Ca^2+^ sensitivity of active SERCA, increasing the Ca^2+^ concentration required for pump cycling. However, PLB does not decrease Ca^2+^ binding to SERCA when ATP is absent, suggesting PLB does not inhibit SERCA Ca^2+^ affinity. The prevailing explanation for these seemingly conflicting results is that PLB slows the Ca^2+^-binding step in the SERCA enzymatic cycle, altering the Ca^2+^ dependence of cycling without affecting the *true* affinity of the Ca^2+^ binding sites. Here, we consider another hypothesis, that measurements of Ca^2+^ binding in the absence of ATP overlook important allosteric effects of nucleotide binding that increase SERCA Ca^2+^ binding affinity. We speculated that PLB inhibits SERCA by reversing this allostery. To test this, we used a fluorescent SERCA biosensor to quantify the Ca^2+^ affinity of non-cycling SERCA in the presence and absence of a non-hydrolyzable ATP-analog, AMPPCP. Nucleotide activation increased SERCA Ca^2+^ affinity, and this effect was reversed by co-expression of PLB. Interestingly, PLB had no effect on Ca^2+^ affinity in the absence of nucleotide. These results reconcile the previous conflicting observations from ATPase assays versus Ca^2+^ binding assays. Moreover, structural analysis of SERCA revealed a novel allosteric pathway connecting the ATP- and Ca^2+^-binding sites. We propose this pathway is interrupted by PLB binding. Thus, PLB reduces the true Ca^2+^ affinity of SERCA by reversing allosteric activation of the pump by ATP.

Significance Statement
The micropeptide phospholamban (PLB) serves a vital role as the “adrenaline trigger” for the heart. PLB inhibits the calcium pump SERCA at rest, and relief of this inhibition increases cardiac performance during exercise. Still, the physical mechanism for PLB inhibition of SERCA remains poorly understood. Here, our results reveal that PLB regulates SERCA by reversing an allosteric effect of ATP that enhances SERCA’s calcium affinity. We mapped a novel allosteric pathway that structurally couples SERCA’s ATP- and calcium-binding sites. PLB binds residues along this pathway to disrupt communication between the ligand-binding sites. The data reveal an allosteric mechanism that may be conserved in other ATP-dependent ion pumps. This allosteric pathway may provide new opportunities for pharmacological targeting of SERCA.

## INTRODUCTION

The sarcoplasmic reticulum Ca^2+^-ATPase (SERCA) sequesters intracellular Ca^2+^ into the lumen of the endoplasmic reticulum (ER) to establish a reservoir for cell signaling. This is a critically important process in all cell types and is energized by ATP hydrolysis and autophosphorylation of the Ca^2+^ pump. Ca^2+^ transport plays a particularly central role in cardiac physiology. The release of Ca^2+^ from the sarcoplasmic reticulum (SR) initiates shortening of the cardiac muscle cell during systole (cardiac contraction). Then, SERCA transport removes Ca^2+^ from the cytosol during diastole (cardiac relaxation) and re-establishes the Ca^2+^ stores in preparation for the next cardiac cycle (*1, 2*). The primary regulator of SERCA function in the heart is phospholamban (PLB), a transmembrane micropeptide that physically interacts with SERCA and inhibits Ca^2+^ transport (*3, 4*). PLB regulation of SERCA is known to be critical for human survival since naturally occurring mutations of PLB that nullify its inhibition are associated with heart failure and premature death by the third decade in carriers (*5*). PLB reduces the Ca^2+^ sensitivity of SERCA during cycling, increasing the Ca^2+^ concentration required for pump turnover (*6, 7*). However, equilibrium measurements of Ca^2+^ binding (in the absence of ATP) have not shown any effect of PLB on the affinity of SERCA for Ca^2+^(*6, 8, 9*). These apparently contradictory results have been reconciled by invoking a kinetic mechanism, that PLB slows the Ca^2+^ binding step of the SERCA enzymatic cycle. This could account for the observed Ca^2+^ desensitization effect of PLB without reducing the true Ca^2+^ affinity of SERCA under equilibrium conditions (*6*).

Alternatively, we speculated that experiments measuring Ca^2+^ binding in the absence of ATP may overlook important allosteric effects of bound nucleotide. In addition to the role of ATP as a source of energy to fuel Ca^2+^ transport, ATP binding to SERCA increases the transporter’s affinity for Ca^2+^. This effect of ATP, referred to as “nucleotide activation”, increases both the rate of Ca^2+^ binding (*10-12*) and the Ca^2+^ affinity of the pump measured using ^45^Ca^2+^ (*13, 14*). Since the effects of nucleotide activation are the opposite of PLB inhibition, we hypothesize that PLB may inhibit SERCA through a mechanism of reversing of nucleotide activation, reducing the true affinity of SERCA for Ca^2+^. Since previous experiments measuring the impact of PLB on Ca^2+^-binding were always performed in the absence of ATP (to prevent enzymatic cycling) (*6, 8, 9*), to our knowledge this possibility has not been investigated. To test this mechanistic hypothesis, we investigated the interplay of ATP binding, Ca^2+^ binding, and PLB binding using a biosensor that reports SERCA conformation through changes in intramolecular fluorescence resonance energy transfer (FRET) (*15*). This biosensor-based assay offers significantly improved sensitivity compared to conventional ^45^Ca^2+^-binding measurements. The results support a new paradigm for the mechanism of regulation of SERCA by PLB.

## RESULTS

### PLB reverses Nucleotide Activation of SERCA Ca^2+^ Affinity by ATP

We previously developed a biosensor called “2-color SERCA” consisting of two fluorescent proteins fused to the A- and N-domains of the cytoplasmic headpiece of SERCA to report its overall conformation by intramolecular FRET (*9, 15, 16*). 2-color SERCA FRET increases in response to increasing Ca^2+^ concentration due to closure of the labeled headpiece domains as the transporter binds Ca^2+^ (*15*) (**Fig. 1A**). Here, we used this 2-color SERCA FRET measurement as an index for relative Ca^2+^ binding to SERCA to study the activating and inhibitory effects of ATP and PLB, respectively, on SERCA Ca^2+^ affinity.

**Figure 1.**
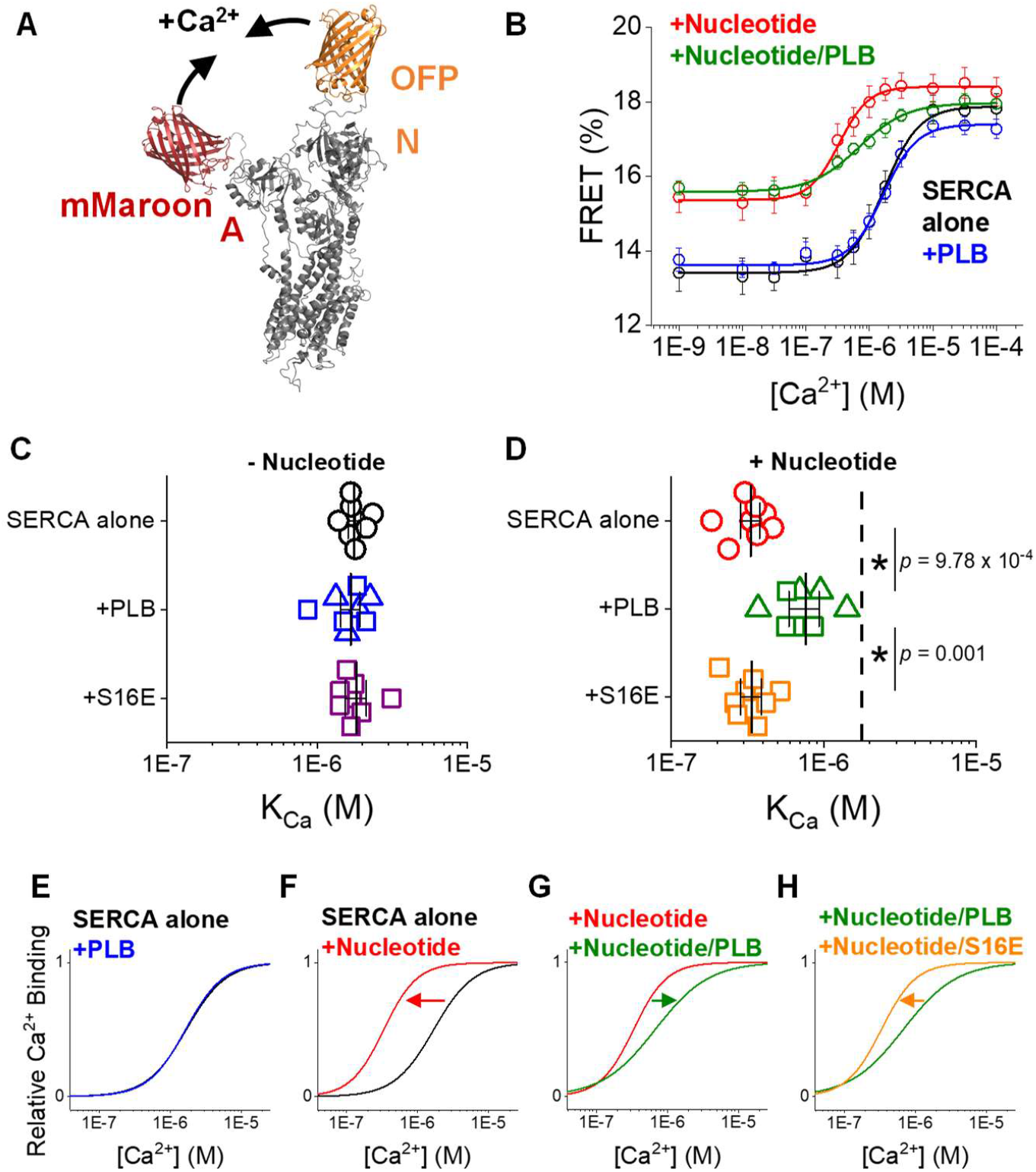
PLB reduces the Ca^2+^ affinity of SERCA by reversing nucleotide activation. **A)** 2-color SERCA biosensor labeled on the A and N domains detects headpiece closure during Ca^2+^ binding by intramolecular FRET. **B)** FRET changes during Ca^2+^ binding measured with SERCA alone (*black*), with AMPPCP (*red*), with PLB co-expressed (*blue*), and with both AMPPCP and PLB (*green*). **C)** Apparent K_Ca_ of SERCA alone (*black*) with co-expression of WT-(*blue*) or S16E-PLB *(purple*) in the absence of nucleotide. **D)** Apparent K_Ca_ of SERCA alone (*red*) with co-expression of WT-(*green*) or S16E-PLB *(orange*) in the presence of nucleotide. SERCA to PLB transfection ratio is indicated by Δ (1:3) or □ (1:5) (*See Fig. S4*). Differences determined by one-way ANOVA with Tukey’s post-hoc (* = *p*<0.05, *n* = 8). See Supplementary Tables S1-4 for complete statistical analysis. **E-H)** Fitted curves from Ca^2+^ binding measurements (as in *B*) normalized to a common minimum and maximum show that PLB inhibition of the relative Ca^2+^ affinity of SERCA is dependent on nucleotide activation and relieved by S16E mutation.

Time-correlated single photon counting (TCSPC) was used to measure changes in the average FRET efficiency of 2-color SERCA expressed in microsomal vesicles, in solutions containing varying concentrations of free [Ca^2+^] (**Fig. 1B**, *black*) (**Fig. S1**, *See methods*). A fit of the data with a Hill function yielded a minimal FRET efficiency for 2-color SERCA at low [Ca^2+^] of 13.5 ± 0.5 % which increased with increasing [Ca^2+^] to a maximum of 17.9 ± 0.4 %. The fit revealed a Ca^2+^ binding constant (K_Ca_) of 1.7 ± 0.1 μM for SERCA alone. This value is consistent with previous measurements of 2-color SERCA Ca^2+^ affinity (*9, 15, 16*). Addition of the non-hydrolyzable ATP analog, AMPPCP (500 μM) shifted the Ca^2+^ binding curve to the left, indicating an increase in Ca^2+^ affinity (K_Ca_ of 333 ± 32 nM, t-test *p* = 6.2E-9) (**Fig. 1B**, *red*). We noted that FRET at low [Ca^2+^] was significantly increased when nucleotide was present to 15.4 ± 0.4 % (*t-test p* = 0.008, **Fig. S2**), consistent with a more compact SERCA headpiece after nucleotide binding (*16, 17*). The increase in SERCA Ca^2+^ affinity with AMPPCP is consistent with previous studies that show that nucleotide binding allosterically increases SERCA’s affinity to subsequently bind Ca^2+^ (nucleotide activation) (*11, 13, 14*).

We investigated the impact of PLB on Ca^2+^ binding to SERCA in the presence and absence of nucleotide using microsomal vesicles co-expressing 2-color SERCA with unlabeled PLB. Since inhibition of SERCA is relieved by PKA phosphorylation of PLB at serine 16, we also tested the effect of co-expressing PLB with a phosphomimetic mutation, S16E (*3, 18-20*) (**Fig. S3**). In the absence of nucleotide, we did not observe any effect on SERCA Ca^2+^ affinity from coexpression of WT- or S16E-PLB (**Fig. 1C and Table S2**). Interestingly, when nucleotide was present, PLB significantly increased the K_Ca_ of SERCA to 782 ± 142 nM (*p* = 9.78E-4) compared to SERCA alone (**Fig. 1D**). This effect of PLB on Ca^2+^ affinity was reversed by S16E mutation, with a significant decrease in the K_Ca_ (337 ± 35 nM, *p* = 0.001) compared to WT-PLB (**Fig. 1D and Table S4**). Taken together, these results suggest that PLB has no effect on Ca^2+^ affinity in the absence of nucleotide (**Fig. 1E**) in agreement with past results (*6, 8, 9*). However, when nucleotide is bound to SERCA, the pump binds Ca^2+^ with much higher affinity (*11-14*), indicated by an appreciable left shift of the concentration dependence of relative Ca^2+^ binding (**Fig. 1F**). Under these biochemical conditions, PLB mediates its primary function of inhibiting SERCA Ca^2+^ affinity by reversing this allosteric activation of the pump by ATP, shifting the curve partially back to the right (**Fig. 1G**). PLB inhibition is relieved when PLB is phosphorylated by PKA. This relief of inhibition is evident from the effect of phosphomimetic S16E mutation of PLB, which shifted the curve back to the left (**Fig. 1H**).

We also evaluated the biosensor response to nucleotide binding. The ATP dependence of the biosensor (K_ATP_ = 10 μM) was compatible with the known ATP-affinity of SERCA (*21*), and ATP binding was not significantly altered by co-expression of PLB (**Fig. S5**).

### MD simulations of SERCA with ATP and PLB

To investigate how nucleotide activation and PLB inhibition impact the structure of SERCA, we performed molecular dynamics (MD) simulations of SERCA with and without ATP bound within the N domain and also in the presence and absence of PLB bound within its regulatory cleft. For simulations with PLB in complex SERCA, the complexes were stable on the microsecond timescale covered by the trajectories. However, in one of the replicate trajectories of ATP-bound SERCA-PLB complex, the ATP became unbound and exited from the nucleotide binding site, and in another replicate the ATP appeared to be loosely attached in its binding pocket Thus, those trajectories should be interpreted with caution.

To evaluate how ATP and PLB impact the structural dynamics of the Ca^2+^ transport sites of SERCA, we analyzed the root mean square fluctuation (RMSF) values for the acidic residues responsible for coordinating Ca^2+^ ions in the binding sites: E309, E771, D800, and E908 (*22, 23*). We noted that the Ca^2+^ gating residue, E309, was more dynamic on this microsecond timescale compared to the other residues, as indicated by higher RMSF values across all trajectories (**Fig. 2A**). Inspection of the Ca^2+^ binding sites revealed that the these residues conform to 2 major geometries during the simulations: (1) a closed conformation where E309 faces the binding pocket and completes the binding sites (**Fig. 2B**) and (2) a more open conformation where E309 is oriented away from the other residues, deforming the Ca^2+^ transport sites (**Fig. 2C**) (*23, 24*). In contrast to E309, E908 displayed the most stability of these residues across trajectories indicated by the lowest RMSF values (**Fig. 2A**). Since E908 is located across from E309 in the Ca^2+^ binding pocket (**Fig. 2B,C**), we monitored the time-dependent change in distance between these residues as an index of the 2 major geometric conformations, with the closed conformation corresponding to a short distance from E908 to E309 (∼15 Å) (**Fig. 2B**, *red dotted line*) and the open conformation characterized by a long distance (∼20 Å) (**Fig. 2C**, *blue dotted line*). **Fig. 2D-G** show the repeated structural transitions between these two conformations over time. We noted that ATP induced a significant ordering of the binding site, with greater sampling of a shorter median E908-E309 distance of 16.2 ± 2.4 Å compared to 15.3 ± 2.1 Å for the APO condition (**Fig. 2D,E** and **S6**). This stabilization of a closed, Ca^2+^-competent conformation is compatible with the observed ATP-dependent increase in Ca^2+^ affinity (Fig. 1B). Interestingly, PLB reversed this nucleotide activation, and the PLB+ATP condition was characterized by greater sampling of a longer median E908-E309 distance of 20.8 ± 1.9 Å (**Fig. 2G** and **S6**). Triangulation of the dynamic E309 residue with E908 and E771 provided additional insight into the apparent disorder-order transition induced by ATP binding. A 2-dimensional density map of the E309-E908 and E309-E771 distances sampled during the simulation shows a wide distribution of distances for the APO condition, with two broad, poorly-defined peaks representing the closed (**Fig. 2H**, *red arrow*) and open conformations (**Fig. 2H**, *blue arrow*). After addition of ATP, there was a marked disorder-to-order transition, resulting in a sharply focused peak at short distances (**Fig. 2I**, *red arrow*). This signifies a shift to a more well-defined, closed conformation. This highly ordered nucleotide-activated conformation of SERCA was abolished by addition of PLB (**Fig. 2J,K**). The 2-D density map shows that addition of PLB shifted the population to an open conformation with a sharp focus at long distances (**Fig. 2K**). Even in the absence of ATP, the presence of PLB resulted in a shift to a longer distance compared to APO (19.1 ± 2.3 Å, **Fig. 2F, J**). Since the APO and PLB-bound conditions show the same low Ca^2+^ affinity in the absence of nucleotide (**Fig. 1E**), we conclude that high affinity Ca^2+^ binding requires both (1) population of the closed conformation of the binding site (**Fig.2E**, *red arrow*) and (2) ordering of the Ca^2+^-binding residues into a sharp focus. This well-ordered arrangement of Ca2+-binding residues results from allosteric activation of SERCA by ATP binding.

**Figure 2.**
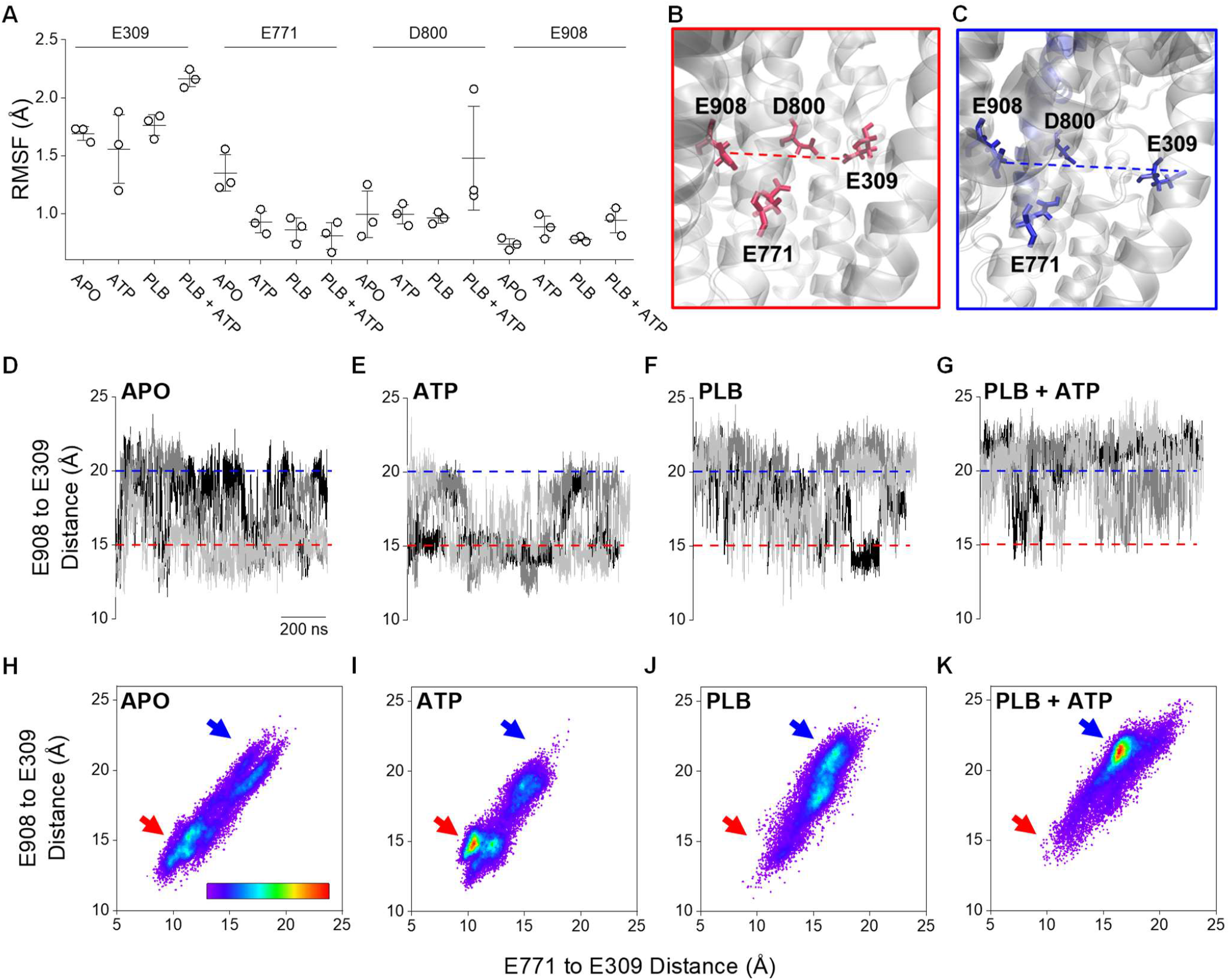
Effects of ATP and PLB on structure of SERCA Ca^2+^ transport sites. **A)** RMSF values for the acidic residues that coordinate Ca^2+^ ions in Ca^2+^ transport sites of SERCA. **B)** Ca^2+^ binding residues shown in a conformation where E309 faces the binding pocket and completes the binding sites **C)** Ca^2+^ binding residues shown in a conformation where E309 is oriented away from the binding pocket, deforming the binding sites. **D-G)** Time-dependent changes in the distance between E908 and E309 for replicates 1 (*black*), 2 (*grey*), and 3 (*light grey*) for each of the simulated conditions. Red and blue dashed lines indicate the distances represented in panels B and C respectively. **H-K)** 2D density maps showing how the distribution of distances occupied by E309 from residues E908 and E771 is affected by the presence of ATP and PLB. Color scale bar represents the density of points ranging from 0 (white) to 21% (red) occupancy. Red and blue arrows indicate the E309 positions represented in panels B and C respectively.

### Allosteric Path Analysis

The changes observed in the structure of the Ca^2+^ binding sites with addition of ATP suggests there indeed exists an allosteric network that couples the structure SERCA binding sites for Ca^2+^ and ATP (**Fig. 1F**). To detect the allosteric path between these sites, we analyzed the molecular dynamics trajectories of APO-SERCA, SERCA+ATP and SERCA+ATP+PLB with GSA tools(*25, 26*). This information theory-based software package is designed to investigate the conformational dynamics of local structures and the functional correlations between local and global motions. In this framework, the protein backbone is represented by a sequence of fragments (f) consisting of four residues, with two consecutive fragments sharing three residues. The local structural states of these fragments over the course of the trajectory were determined using a structural alphabet (*27*), and we calculated the mutual information (MI) between fragments as a measure of the correlation between the local states of the fragments. A network is constructed for the protein based on the MI for all the pair of fragments. The fragments associated with ATP binding and autophosphorylation sites act as one set of endpoints at one end of the allosteric network, and those fragments related to the Ca^2+^ binding site act as the other set of endpoints at the other end. We do not get an allosteric path between these endpoints for the APO-SERCA due to high-weight (low MI) connections between the fragments. The addition of ATP significantly improves the coupling, and we obtain an allosteric path passing through the following fragments: *f*560 → *f*556 → *f*362 → *f*124 → *f*115 → *f*917 → *f*771 (**Fig. 3**). Of note, the path passes through *f*115 of TM2 which is part of the PLB regulatory cleft in the transmembrane domain of SERCA, suggesting that PLB plays a role in the modulation of allosteric signal from the ATP binding site to the Ca^2+^ binding site. The residues corresponding to each fragment in this path are shown in **Table 1**. So, the allosteric path only exists when ATP is bound to SERCA, and not in its APO state. For SERCA+ATP+PLB, we do not obtain any allosteric path as the Ca^2+^ and ATP binding sites are either not connected in the network or connected with edges having higher weights (low MI). This indicates that the addition of PLB along with ATP bound to SERCA inhibits the allosteric communication between ATP and Ca^2+^ binding sites and reverses the nucleotide activation.

**Figure 3.**
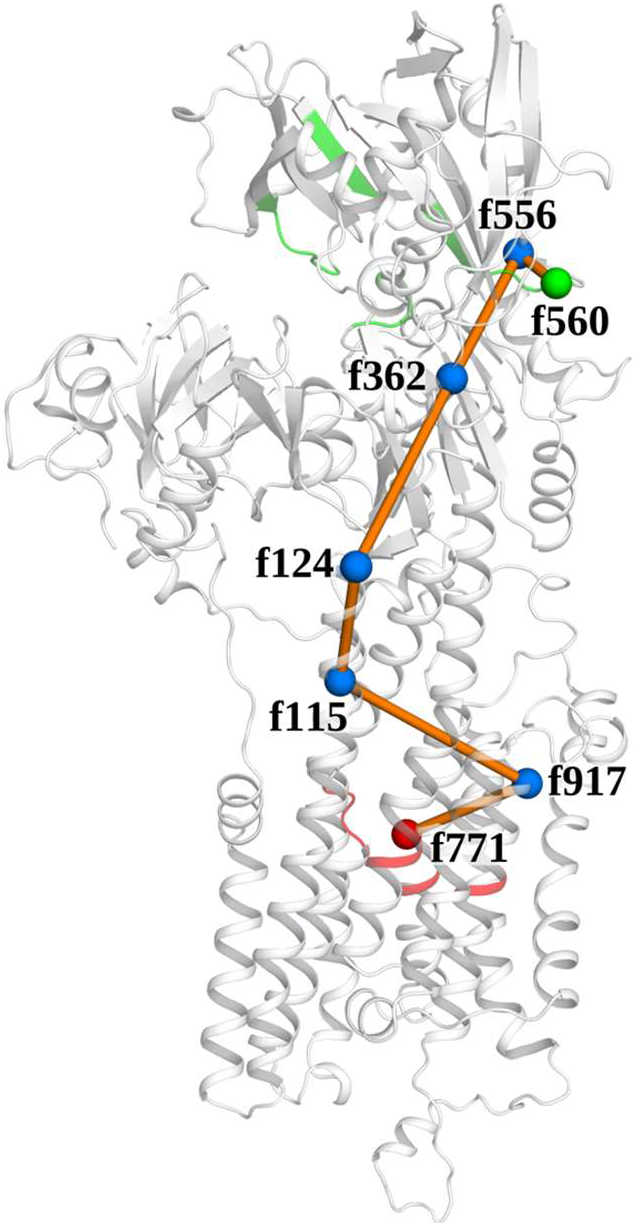
Mapping the allosteric pathway for nucleotide activation of SERCA. Analysis of MD simulations of SERCA ± ATP/PLB with GSA tools revealed an allosteric pathway (*orange line + blue dots*) that coupled the structural dynamics of the ATP-(*green*) and Ca^2+^-binding sites (*red*) of SERCA in trajectories with ATP bound. PLB binds with residues in fragment 115 (*f115*) in TM2 and disrupted the allosteric coupling of this pathway.

**Table 1.**
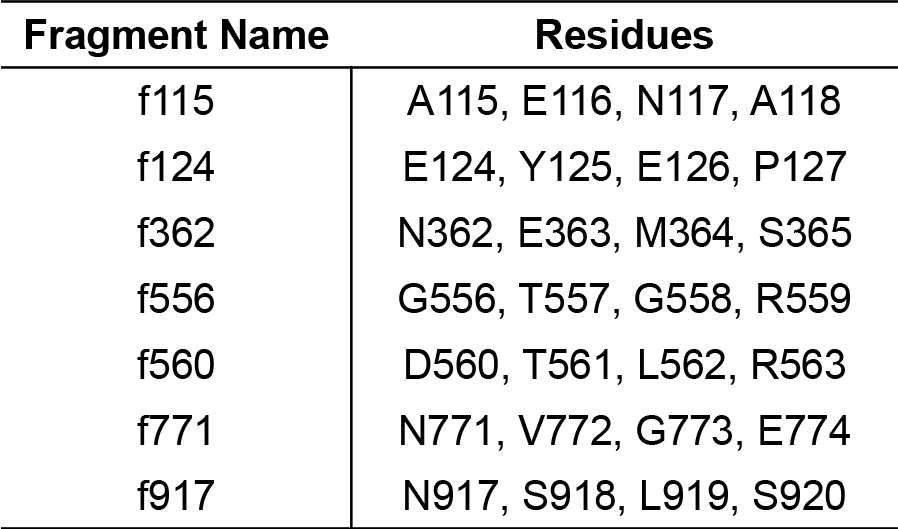
Residues identified in the allosteric path coupling the ATP and Ca^2+^ binding sites.

| Fragment Name | Residues |
| --- | --- |
| f115 | A115, E116, N117, A118 |
| f124 | E124, Y125, E126, P127 |
| f362 | N362, E363, M364, S365 |
| f556 | G556, T557, G558, R559 |
| f560 | D560, T561, L562, R563 |
| f771 | N771, V772, G773, E774 |
| f917 | N917, S918, L919, S920 |

## DISCUSSION

In previous studies, we (*28-30*) and others (*6, 31-33*) have investigated SERCA regulation by PLB under the premise that PLB reduced the kinetics of Ca^2+^ binding to SERCA, altering the pump’s “apparent” Ca^2+^ affinity without changing the actual affinity of the Ca^2+^ binding sites. This perspective was based on Ca^+2^-binding experiments performed in the absence of ATP (*6, 8, 9*), a condition routinely used to stop SERCA enzymatic cycling for equilibrium measurements. In the present study, we prevented enzymatic cycling of the pump using a non-hydrolyzable analog of ATP, which enabled quantification of Ca^2+-^binding to the nucleotide-bound SERCA under equilibrium conditions. These experiments also exploited a fluorescent biosensor, “2-color SERCA”, that reports the Ca^2+^-dependent conformation change in the SERCA headpiece with a change in FRET (*15*). This assay offers an advantage of 2-3 orders of magnitude improved sensitivity compared to conventional quantification of ^45^Ca^2+^ binding to cardiac SR (*6*). The results recapitulate several key observations from previous studies. We found that the presence of nucleotide greatly increased the Ca^2+^ sensitivity of SERCA, in agreement with past results (*10-14*). This phenomenon is referred to as “nucleotide activation” (**Fig. 1F**). We also reproduced the observation that there is no effect of PLB on equilibrium Ca^2+^ binding to SERCA in the absence of nucleotide (**Fig. 1E**) (*6, 8, 9*). However, PLB *does* decrease SERCA’s equilibrium Ca^2+^ binding in the presence of non-hydrolyzable nucleotide (**Fig. 1G**). Others have previously speculated that nucleotide might be needed to detect the inhibitory effect of PLB (*32, 34*), but to our knowledge, this is the first observation that PLB decreases the true affinity of the Ca^2+^ binding sites of non-cycling SERCA. The data suggests that the mechanism of action of PLB involves modulating the allosteric connection between the ATP- and Ca^2+^-binding sites.

MD simulations provided insight into this allosteric mechanism. Of particular interest were the dynamic motions we observed in glutamine 309 (**Fig. 2A**). This residue has well-established roles in both gating the entry of Ca^2+^ ions into the binding cavity (*23, 24*) and allosterically coupling the structure of the TM and headpiece domains of SERCA (*35-37*). In the APO condition, we observed this residue moving in and out of the Ca^2+^ binding pocket (**Fig. 2H**), consistent with crystal structures that have shown E309 in either position. E309 faces the binding pocket in Ca^2+^ bound states of SERCA (*38, 39*) but faces away when thapsigargin is bound and inhibits Ca^2+^ binding (*23*). ATP and PLB both impacted the Ca^2+^ binding sites by affecting equilibrium position of this residue. In simulations with ATP bound, our results showed E309 more frequently occupied a well-defined position in the closed conformation (**Fig 2B**,**I**), suggesting that ATP binding to SERCA allosterically primes the transporter for Ca^2+^ binding. This is in agreement with past biochemical studies that found E309 contributes to the enhancement of Ca^2+^ binding and occlusion by adenosine nucleotides (*24*). Interestingly, simulations with PLB bound in complex with SERCA showed the opposite trend, with the PLB+ATP condition exhibiting a strong preference for occupying the more open state of E309 where the Ca^2+^ binding site is less formed (**Fig. 2C**,**K**). This suggests that the PLB interaction with SERCA disrupts the allosteric communication between the ATP and Ca^2+^ binding sites and stabilizes the pump in a less competent conformation for binding Ca^2+^.

The allosteric network analysis provides additional insight into the results of the biochemical measurements and MD simulations. This analysis revealed a novel allosteric pathway that couples ATP- and Ca^2+^ binding sites of SERCA when ATP is bound. Interestingly, this route passes through the PLB binding cleft residues on TM2. The presence of PLB disrupts the structural coupling between the remote ligand-binding sites, most likely by altering the conformations of the residues in the PLB-binding regulatory site: A115, E116, N117, and A118. It was noteworthy that E771 was the end point of the predicted pathway because we have previously shown that this residue plays a key role in PLB-mediated modulation of SERCA (*29*). Since this allosteric pathway is utilized by both nucleotide activation and PLB inhibition of the pump, the structural elements along this pathway may present new targets for pharmacological stimulation or inhibition of SERCA.

The present study reveals insight into the allosteric regulation of SERCA by ATP and PLB. Additional context for these results is provided by our previous study (*28*), in which we showed that the PLB-SERCA regulatory complex is most stable when the Ca^2+^ pump is in its ATP-bound conformation (*32, 40-42*). Interestingly, this ATP-dependent increase in PLB-SERCA affinity was reversed upon Ca^2+^ binding (*28*). **Figure 4** summarizes the allosteric relationships between SERCA binding sites for ATP, Ca^2+^, and PLB. ATP binding to its active site in the N-domain allosterically activates the Ca^2+^ binding sites in the transmembrane domain, increasing SERCA Ca^2+^ affinity (**Fig. 4**, *green*). ATP binding also causes PLB to interact more avidly with its transmembrane regulatory cleft (**Fig. 4**, *orange*). Here, we demonstrated that PLB inhibits SERCA by disrupting the allosteric coupling of the ATP- and Ca^2+^-binding sites (**Fig. 4**, *blue*), reversing nucleotide activation of SERCA Ca^2+^ binding. Counter regulation by Ca^2+^ occurs as the affinity of the PLB-SERCA regulatory complex with ATP is reduced after Ca^2+^ is bound (**Fig. 4**, *red*). Overall, the results provide new insight into a novel allosteric pathway connecting the ATP binding site with distant Ca^2+^ binding sites and reveal that PLB disrupts this path.

**Figure 4.**
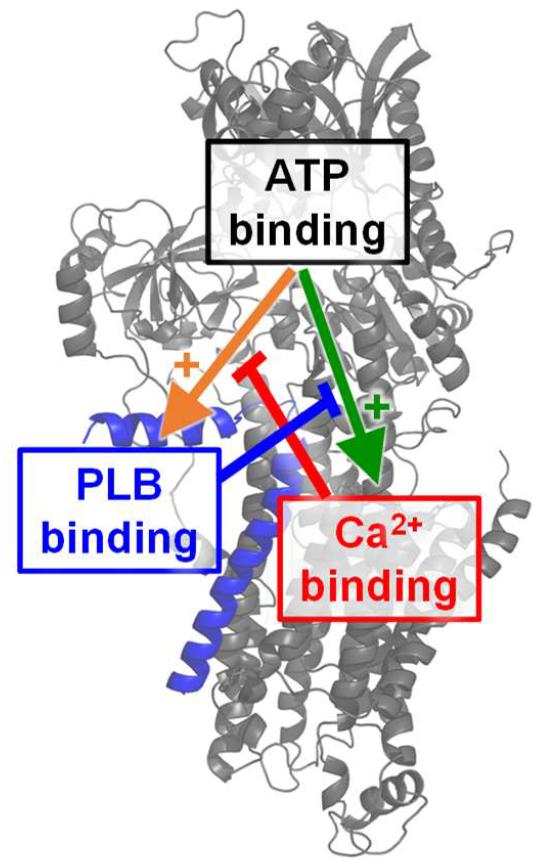
A schematic diagram of the allosteric interplay between ATP, PLB, and Ca^2+^ binding to SERCA.

## METHODS

### Plasmid Constructs

The engineering and functional characterization of our canine 2-color SERCA2a has been previously described (*15, 16, 43, 44*). Here, we used a version of this construct labeled with mMaroon1 (mMaroon) on the N-terminus, labeling the A domain, and an mCyRFP1 (OFP) intersequence tag inserted before residue 509 on the N domain of SERCA. The OFP and mMaroon donor/acceptor FRET pair has a Förster distance (*R*_*0*_) of 63.34 Å. Our lab has previously demonstrated that the fusion of one or two fluorescent proteins to SERCA did not alter normal Ca^2+^ transport function (*15, 16, 44*). Furthermore, PLB fused to another tag was able to normally regulate SERA function (*15*). Therefore, we believe fluorescent proteins are benign for the normal function of SERCA or PLB.

### Molecular biology and cell culture

HEK-293T cells were cultured in Dulbecco’s modified Eagle’s medium (DMEM) supplemented with 10% fetal bovine serum (ThermoScientific). Cells were cultured on 150 mm^2^ dishes and transiently transfected using the Lipofectamine 3000 transfection kit (Invitrogen) with either (*1*) 90 μg of 2-color SERCA plasmid DNA alone or (*2*) 50 μg of 2-color SERCA plasmid DNA and unlabeled PLB plasmid DNA supplemented at a 1:3 or 1:5 SERCA to PLB ratio (150 or 250 μg unlabeled PLB, respectively).

### HEK-293T cell microsomal membrane preparation

ER microsomal membranes expressing 2-color SERCA were isolated as previously described (*16*). Briefly, roughly 48 hours post-transfection, cells expressing 2-color SERCA were washed with PBS, harvested by scraping in 20 mL of an ice-cold homogenizing solution containing 10 mM Tris-HCl pH 7.5, 0.5 mM MgCl_2_, and an EDTA free protease inhibitor cocktail, and pelleted by centrifugation at 1000 x g for 10 min at 4 °C. Cell pellets were resuspended in 5 mL of cold homogenizing solution and disrupted by 10 strokes in a Potter-Elvehjem homogenizer. Cell homogenates were then supplemented with 5 mL of ice-cold sucrose solution (100mM MOPS pH 7.0, 500 mM sucrose, and an EDTA free protease inhibitor cocktail) and passed throusgh a 27-gauge needle 10 times. Cell homogenates were then centrifuged at 1000 x g for 10 min at 4 °C. The supernatants were collected and centrifuged at 126,000 x g for 30 min at 4 °C. High speed membrane pellets were resuspended in a 1:1 mixture of homogenizing and sucrose solutions, disrupted by 10 strokes in a Potter-Elvehjem homogenizer, and passed through a 27-gauge needle 10 times. A Pierce BCA assay kit (ThermoScientific) was used to determine the protein concentration of membrane preparations.

### Time-correlated single-photon counting

Fluorescence lifetime measurements were obtained from microsomal membrane preparations from HEK-293T cells expressing 2-color SERCA labeled with mMaroon1- and mCyRFP1-labeled on the A and N domains respectively, with or without unlabeled PLB co-expressed. Membranes were diluted at a 1:10 ratio in a solution containing 100 mM KCl, 5 mM MgCl_2_, 10 mM imidazole, 2 mM EGTA, and varying concentrations of CaCl_2_. For the nucleotide bound condition, 500 μM AMPPCP was added to solutions. The mCyRFP1 donor was excited with a supercontinuum laser (FIANIUM) filtered through a 482/18 nm bandpass filter. Emitted fluorescence from the sample was detected through a 1.2 N.A. water immersion objective and transmitted through a 593/40 nm bandpass filter to a PMA Hybrid series detector (PicoQuant). Light from the detector was quantified by a HydraHarp 400 single-photon counting module at 16 ps resolution. TCSPC histograms were obtained over 60 s acquisition period for each condition. In control experiments with singly-labeled mCyRFP-SERCA, the donor alone gave a single-exponential decay with a *τ*_*D*_ of 3.52 ns. TCSPC histograms from 2-color SERCA samples were fit with a 2-exponential decay function, which was used to derive the average lifetime for the two 2-color SERCA populations (*τ*_*DA*_). FRET efficiencies were calculated for each sample according to the relationship 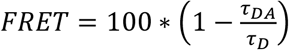 (*45*) and plotted as a function of Ca^2+^ concentration. The data were well described by a Hill function of the form 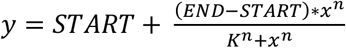, where *START* is the minimal FRET efficiency at low Ca^2+^, *END* is the maximum FRET efficiency at high Ca^2+^, *n* is the Hill coefficient, and *K* is the Ca^2+^ binding constant (K_Ca_). Data from 8 independent experiments from a minimum of 4 microsomal preparations were global fit to obtain a single best Hill coefficient with independent K_Ca_ values for each experimental condition. For Ca^2+^ binding measurements with microsomes coexpressing 2-color SERCA and unlabeled WT-PLB, we evaluated 3:1 and 5:1 PLB to SERCA expression ratios. We did not detect a significant difference in Ca^2+^ affinity between these samples, but observed improved cooperativity of Ca^2+^ binding with a higher PLB to SERCA expression ratio (**Fig. S4**) (*46*).

### Preparation of the systems

We used the crystal structure of SERCA (PDB 3w5a) to simulate the SERCA apo and SERCA–ATP complexes. To simulate the SERCA–PLB systems, we used the atomic model of the full-length complex previously reported by us (*47*). To model the ATP-bound structure, we docked ATP onto the nucleotide-binding pocket of SERCA using the CB-DOCK program (*48*). We adjusted the pK_a_ of other ionizable residues to a pH value of ∼7.2 using PROPKA (*49, 50*). The complexes were embedded in a 120×120 Å bilayer of POPC lipids. The initial system was solvated using TIP3P water molecules with a margin of 20 Å in the z-axis between the edges of the periodic box and the cytosolic and luminal domains of SERCA, respectively. K^+^, and Cl^-^ ions were added to neutralize the system and to produce a KCl concentration of ∼100 mM. Preparation of the systems was done using the CHARMM-GUI web interface.(*51*)

### Molecular dynamics simulations

We performed molecular simulations with AMBER20 on Tesla V100 GPUs (*52*) using the AMBER ff19SB force field.(*53*) We maintained a temperature of 310 K with a Langevin thermostat and a pressure of 1.0 bar with the Monte Carlo barostat. We used the SHAKE algorithm to constrain all bonds involving hydrogens and allow a time step of 2 fs. We first performed 5000 steps of steepest-descent energy minimization followed by equilibration using two 25-ps MD simulations using a canonical ensemble (NVT), one 25-ps MD simulation using an isothermal–isobaric ensemble (NPT), and two 500-ps MD simulations using the NPT ensemble. The equilibrated systems were used as a starting point to perform the production MD simulations.

### Information-theoretic analysis of allosteric paths in SERCA

The simulations were done for four configurations of SERCA: APO-SERCA, SERCA+ATP, SERCA+PLB and SERCA+ATP+PLB. The trajectories for the four configurations were analyzed to detect the allosteric path between ATP binding/autophosphorylation site and Ca^2+^ binding site using GSA tools to determine conformational dynamics and functional correlations between local and global motions (*25, 26*). The protein backbone was represented by overlapping fragments consisting of four residues, and the local structural states of these fragments over the course of the trajectory were determined using a structural alphabet, as previously described (*27*). The mutual information (MI) between fragments was calculated to determine the correlation between the local states of the fragments. The normalized mutual information between two fragments *i* and *j*, represented by 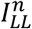 is,

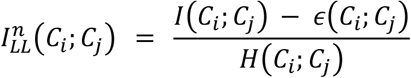

Here, *C*_*i*_ and *C*_*j*_ : columns *i* and *j* in the string alignment. *I(C*_*i*_ ; *C*_*j*_ ): mutual information, *H(C*_*i*_, *C*_*j*_): joint entropy of two fragments. *ε(C* ; *C*_*j*_ ) : error term arising from finite size error. Using the 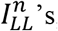, we constructed a network with different fragments represented as nodes which are connected by the edges. The MI between the two fragments determines the weight of an edge between each node pair *(i, j)*, and is given by the relation,

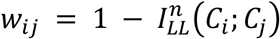

In this analysis, an appropriate value of the distance cutoff and MI cutoff must be assigned. The distance cutoff sets the maximum physical distance between the first Cα atoms of the fragment pair for which an edge connection between the fragments can exist. A relevant value of cutoff for the MI between the nodes 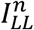 have also to be set so that the edges with high weights (or low 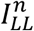) can be neglected. The network thus constructed for a protein is sensitive to these cutoffs. In our analysis, we set the distance cutoff to be 30 Å and the 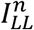 cutoff was taken to be 33% of the maximum value of MI between any pairs of fragments for the four configurations and is equal to 0.133. So, a pair of nodes will have an edge 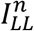 if the is in the top 67 % of all the 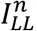 between different pairs of fragments and within the distance cutoff of 30 Å. Ideally, one should take the MI between the fragments 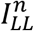 to be greater than 50%. We set a lower cutoff because we did not get a path between two binding sites for a cutoff greater than 50% of SERCA-ATP. The endpoints of the possible allosteric network are taken as the fragments associated with Site A (ATP binding site and auto-phosphorylation site) and Site B (Ca^2+^ binding site.) We ascertain whether the paths exist between sites A and B and then determine the shortest paths (i.e., paths with the lowest cost) using Dijkstra’s algorithm for the four cases. A relevant quantity of interest here is the node eigenvector centrality which measures the relative importance of each node in the network. The highest-centrality nodes, which have a large number of connections and are likely to be involved in allosteric signal transmission, were determined for each case. A normalized mutual information 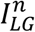 between the local states of the fragment and the global collective motion is also calculated. This identifies the fragments which are likely to be associated with the collective global motion of the protein and hence are important in determining the allosteric path (*25*).

## ACKNOWLEDGEMENTS

The authors would like to thank Mike Autry, Zhenhui Chen, and Audrey Deyawe Kongmeneck for helpful discussions. This investigation was supported by the National Institutes of Health (NIH): Ruth L. Kirschstein Predoctoral Individual National Research Service Award (NRSA) F31HL165900-01 from the National Heart, Lung, and Blood Institute (NHLBI) to S.R.C; R01HL092321 and R01HL143816 from NHLBI to S. L.R.; R01GM120142 from the National Institute of General Medical Sciences (NIGMS) to L.M.E.F. KB is supported by by a Villum foundation Grant number 35888. HK is supported by a Lundbeckfonden Ascending Investigator #R344-2020-1023.

## Notes

### Competing Interest Statement

The authors have declared no competing interest.

