## Supplementary material for "Phospholamban Inhibits the Cardiac Calcium Pump Through Reversing the Allosteric Enhancement of Calcium Affinity by ATP": Fig. S1

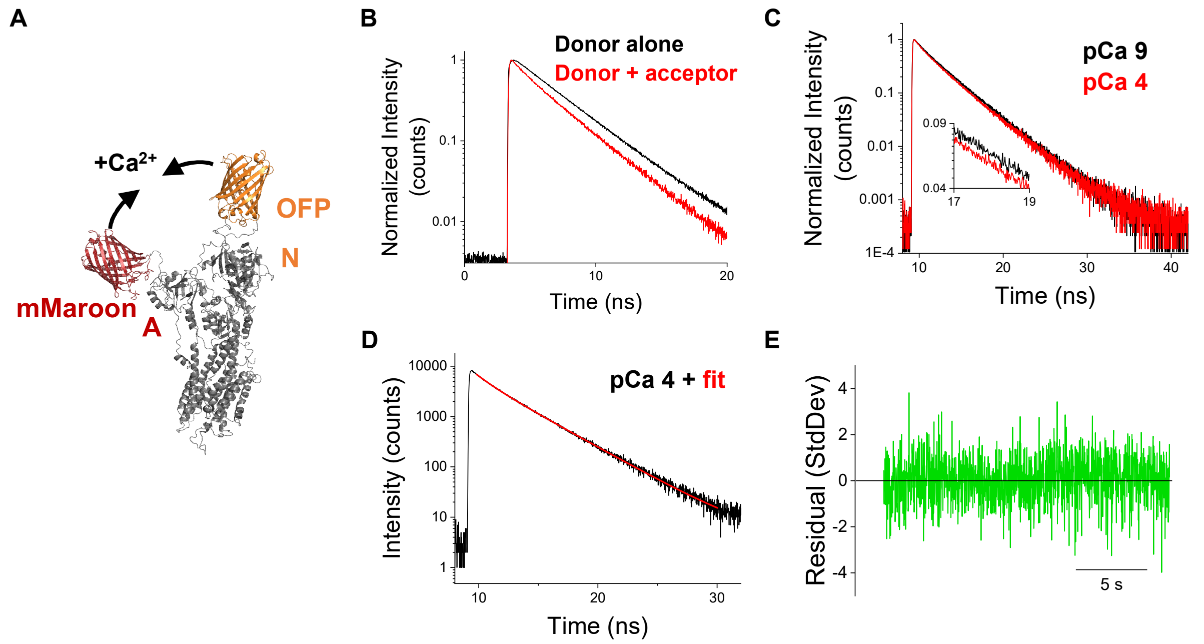


**Supplementary Figure S1. A)** 2-color SERCA biosensor labeled on the A and N domains undergoes headpiece closure during $Ca^{2+}$ binding by intramolecular FRET. **B)** TCSPC measures how the fluorescence lifetime of the OFP donor tag on the N domain of SERCA decreases due to FRET in the presence of an mMaroon acceptor (*red*) on the A domain compared to the lifetime of SERCA tagged with OFP alone (*black*). **C)** The fluorescence lifetime of the OFP donor in 2-color SERCA is further decreased when $Ca^{2+}$ is increased from pCa 9 to pCa 4, due closure of the labeled headpieces (as in panel A). The inset data are zoomed in two make the difference more appreciable. **D)** A representative fit (*red*) of the OFP donor lifetime (*black*) using a two exponential decay function. **E)** A residual of a two-exponential decay fit of the OFP donor lifetime of 2-color SERCA shows that the data are well described by this two species model.
