## Supplementary material for "Phospholamban Inhibits the Cardiac Calcium Pump Through Reversing the Allosteric Enhancement of Calcium Affinity by ATP": Fig. S5

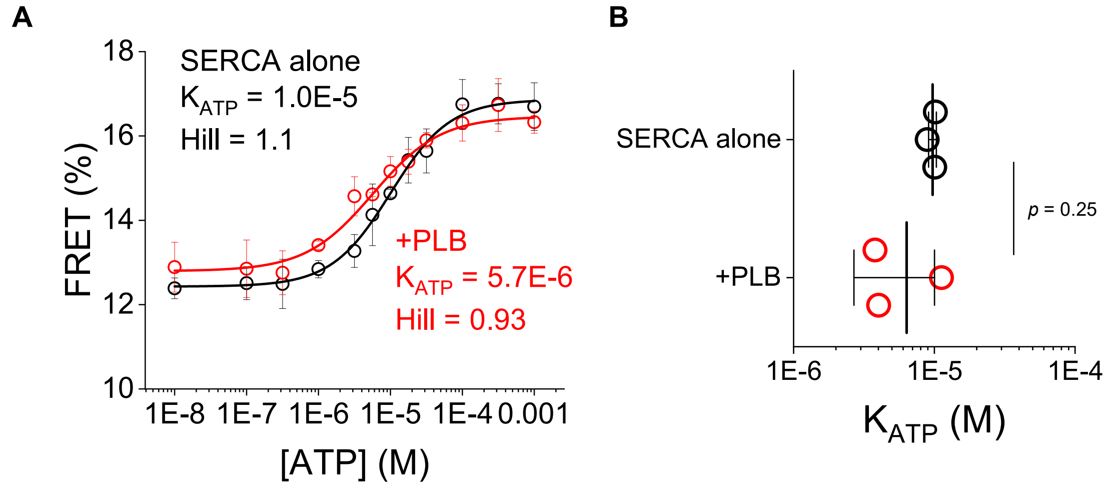


**Supplementary Figure S5. A)** 2-color SERCA FRET changes during ATP binding measured in samples expressing SERCA alone (*black*) or coexpressing WT-PLB at a 1:5 SERCA to PLB ratio (*red*). **B)** Values for the apparent ATP binding constant ($K_{ATP}$) of 2-color SERCA determined from 3 independent experiments. Differences determined by student's t-test (*= $p<0.05$).
