## Supplementary material for "Phospholamban Inhibits the Cardiac Calcium Pump Through Reversing the Allosteric Enhancement of Calcium Affinity by ATP": Fig. S6

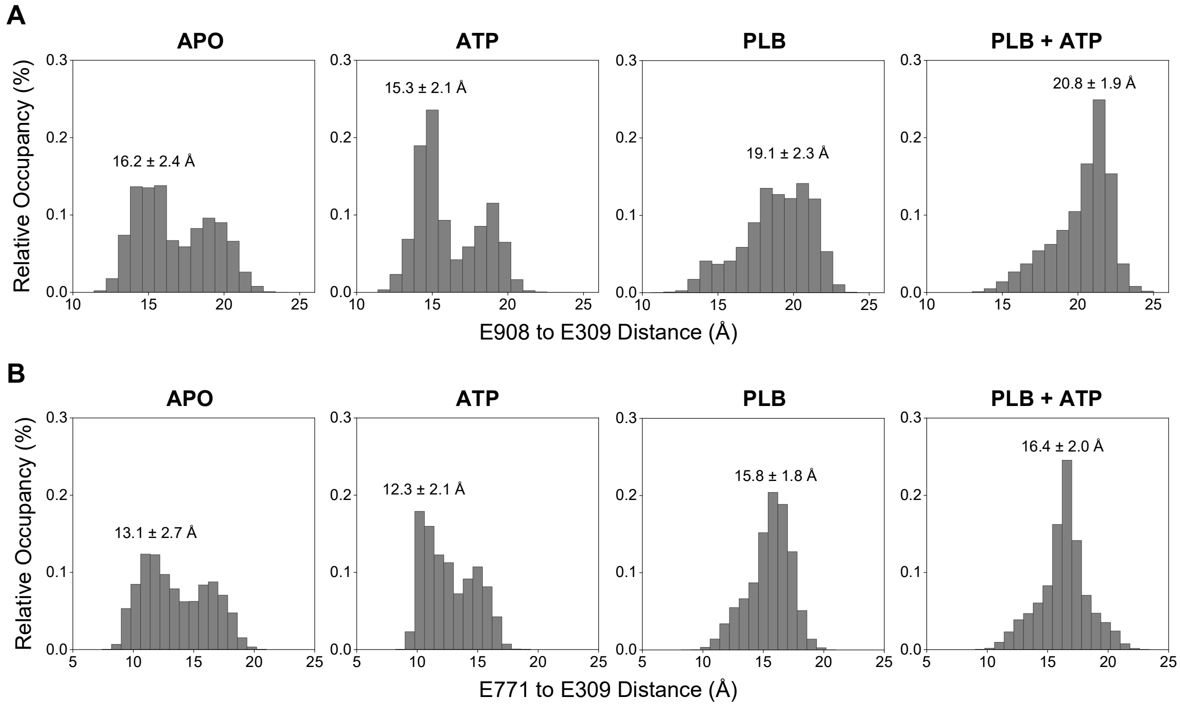


**Supplementary Figure S6. A)** Relative occupancy of the distances between SERCA $Ca^{2+}$ binding residues E908 and E309 based on 3 independent trajectories. **B)** Relative occupancy of the distances between SERCA $Ca^{2+}$ binding residues E771 and E309 based on 3 independent trajectories. Overlayed values are the median and standard error for each data set.
