## Supplementary material for "Phospholamban Inhibits the Cardiac Calcium Pump Through Reversing the Allosteric Enhancement of Calcium Affinity by ATP": Fig. S2

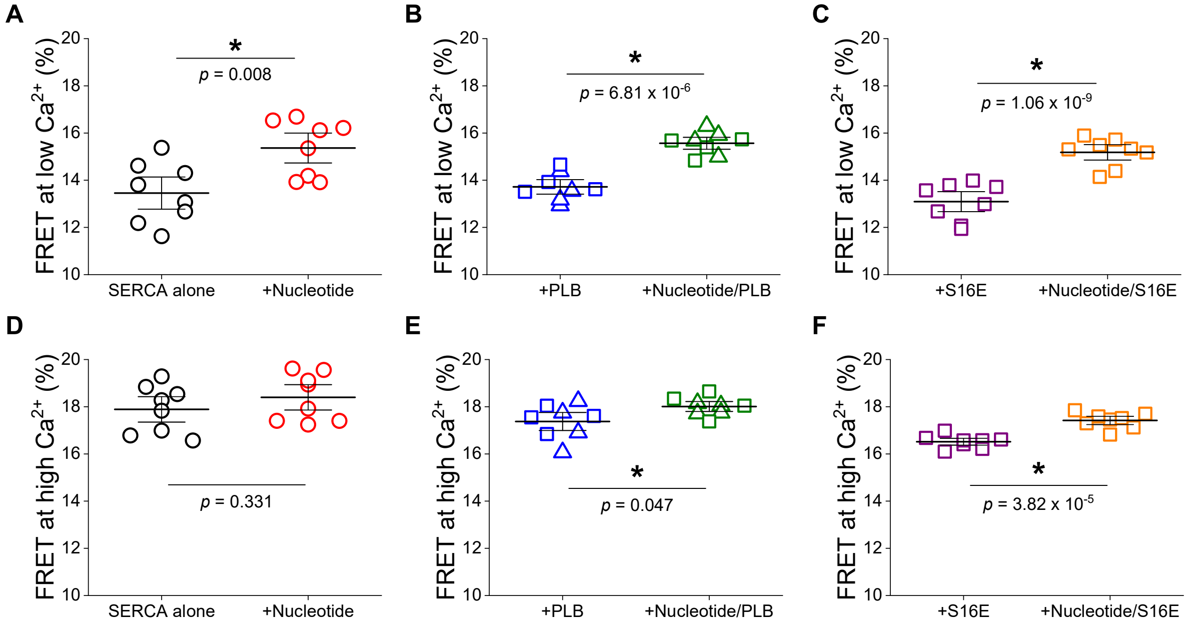


**Supplementary Figure S2.** The FRET efficiency of 2-color SERCA under low and high $Ca^{2+}$ conditions as determined by fitting the average FRET of the biosensor in a range of $Ca^{2+}$ concentrations with a Hill equation. **A-C)** FRET efficiency of 2-color SERCA at low $Ca^{2+}$ was significantly increased by nucleotide activation in all samples. **D-F)** FRET efficiency of 2-color SERCA at high $Ca^{2+}$ was significantly increased only in samples coexpressing WT- or S16E-PLB. Differences determined by student's t-test (*= $p$<0.05)
