## Supplementary material for "Phospholamban Inhibits the Cardiac Calcium Pump Through Reversing the Allosteric Enhancement of Calcium Affinity by ATP": Fig. S3

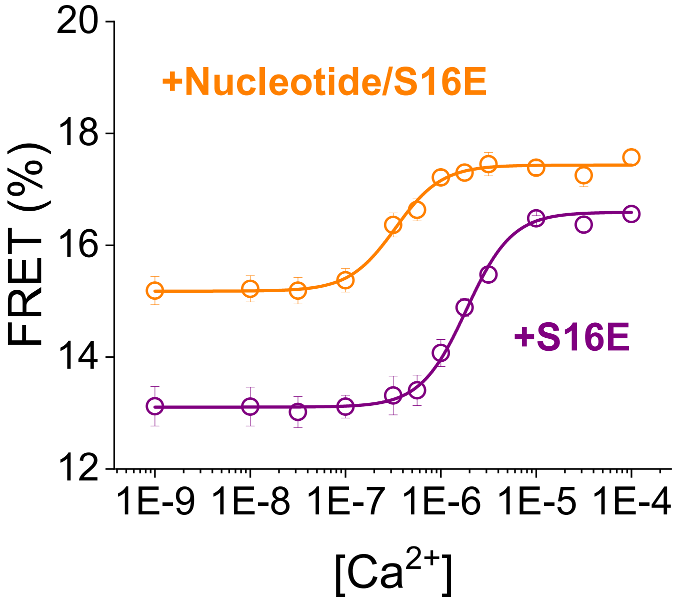


**Supplementary Figure S3.** 2-color SERCA FRET changes during $Ca^{2+}$ binding measured in samples coexpressing SERCA and S16E-PLB (1:5 SERCA to PLB ratio) in the presence (*orange*) and absence of AMPPCP (*purple*).
