## Supplementary material for "Phospholamban Inhibits the Cardiac Calcium Pump Through Reversing the Allosteric Enhancement of Calcium Affinity by ATP": Fig. S4

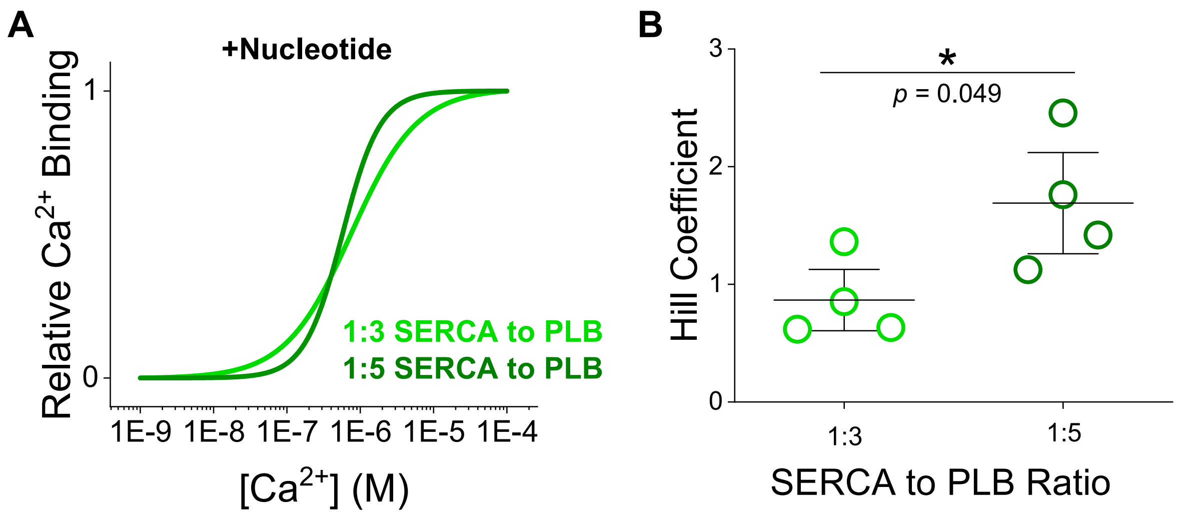


**Supplementary Figure S4. A)** Normalized fitted curves for 2-color SERCA FRET changes during $Ca^{2+}$ binding in the presence of AMPPCP for samples coexpressing WT-PLB at a 1:3 (*light green*) or a 1:5 (*dark green*) SERCA to PLB ratio. Data are based on fitting the mean and SE of 4 independent measurements. **B)** Hill coefficient values for changes in 2-color SERCA FRET during $Ca^{2+}$ binding for samples coexpressing WT-PLB at a 1:3 (*light green*) or a 1:5 (*dark green*) SERCA to PLB ratio. Hill coefficient values were determined by global fitting the data from 4 independent experiments to share a common $K_{Ca}$ value. Differences determined by student's t-test (*= $p$<0.05).
