## Supplementary material for "Phospholamban Inhibits the Cardiac Calcium Pump Through Reversing the Allosteric Enhancement of Calcium Affinity by ATP": Table S3

**Supplementary Table S3.** Apparent $Ca^{2+}$ binding constants derived from intramolecular FRET measurements of 2-color SERCA alone and with WT- or S16E-PLB in the presence of AMPPCP.

| **Apparent $K_{Ca}$ +Nucleotide (Mean ± SEM)** | |
|---|---|
| Condition | $K_{Ca}$ (μM) |
| SERCA alone | 0.33 ± 0.03 |
| PLB | 0.78 ± 0.14 |
| S16E | 0.34 ± 0.04 |
