## Supplementary material for "Phospholamban Inhibits the Cardiac Calcium Pump Through Reversing the Allosteric Enhancement of Calcium Affinity by ATP": Table S4

**Supplementary Table S4.** *P* values comparing differences in apparent $Ca^{2+}$ binding constants of SERCA alone and with WT- or S16E-PLB in the presence of AMPPCP. These values were determined by one-way ANOVA with Tukey's *post-hoc* test (* = $p<0.05$).

| Apparent $K_{Ca}$ +Nucleotide<br>*p* values from 1-way ANOVA with Tukey's post-hoc | SERCA alone | PLB |
|---|---|---|
| S16E | 0.999 | **0.001*** |
| PLB | **9.78 x $10^{-4}$*** | |
