## Supplementary material for "Phospholamban Inhibits the Cardiac Calcium Pump Through Reversing the Allosteric Enhancement of Calcium Affinity by ATP": Table S1

**Supplementary Table S1.** Apparent $Ca^{2+}$ binding constants derived from intramolecular FRET measurements of 2-color SERCA alone and with WT- or S16E-PLB in the absence of nucleotide.

| Apparent $K_{Ca}$ in the Absence of Nucleotide (Mean ± SEM) | |
|---|---|
| Condition | $K_{Ca}$ (μM) |
| SERCA alone | 1.77 ± 0.11 |
| PLB | 1.68 ± 0.16 |
| S16E | 1.84 ± 0.20 |
