## Supplementary material for "Phospholamban Inhibits the Cardiac Calcium Pump Through Reversing the Allosteric Enhancement of Calcium Affinity by ATP": Table S2

**Supplementary Table S2.** *P* values comparing differences in apparent $Ca^{2+}$ binding constants of SERCA alone and with WT- or S16E-PLB in the absence of nucleotide. These values were determined by one-way ANOVA with Tukey's *post-hoc* test (* = $p<0.05$).

| Apparent $K_{Ca}$ in the Absence of Nucleotide<br>*p* values from 1-way ANOVA with Tukey's post-hoc | | |
|---|---|---|
| | SERCA alone | PLB |
| S16E | 0.959 | 0.762 |
| PLB | 0.903 | |
